# taxmyPHAGE: Automated taxonomy of dsDNA phage genomes at the genus and species level

**DOI:** 10.1101/2024.08.09.606593

**Authors:** Andrew Millard, Remi Denise, Maria Lestido, Moi Nicholas, Deven Turner, Dann Turner, Thomas Sicheritz-Pontén

## Abstract

**Background:** Bacteriophages are now classified into genera and species based on genomic similarity. To identify new species and genera requires comparison against all phage genomes currently defined by the International Committee on the Taxonomy of Viruses. With increasing amounts of phage genomic data there is a clear need to fully automate the classification of phages into existing and new genera and species.

**Materials and Methods:** We created a MASH database for rapid searching of closely related phages, to then be compared with BLASTn to compute intergenomic similarity between bacteriophages (https://github.com/amillard/tax_myPHAGE/tree/main).

**Results:** Taxmyphage is provided as a conda repository and via a user-friendly website, that allows rapid classification of phages into existing taxa at the genus and species level.

**Conclusions:** Taxmyphage allows the rapid and accurate classification of phage genomes into existing genera and species, at a scale that is compatible with existing sequencing output.

## Introduction

Bacteriophages are viruses that specifically infect bacteria, are ubiquitous and some of the most abundant biological entities on the planet. Unlike their bacterial hosts they do not have to maintain their genome as dsDNA, with some bacteriophages utilising ssRNA or ssDNA as their genetic material. Additionally, we now know bacteriophage genomes span a large size range from ∼3.3 kbp (1) of ssRNA bacteriophages to greater than 700 kbp (2,3).

The classification of bacteriophages into hierarchical groups based on their evolutionary relationships (i.e. taxonomy) and regulated naming of such groups (i.e.nomenclature) has evolved considerably since their first discovery in the early 20th century. Viral taxonomy is overseen by the International Committee on Taxonomy of Viruses (ICTV). Since its establishment in 1966, the committee has been responsible for developing, refining, and maintaining a universal system of virus taxonomy (4). Given their small size and absence of accessible sequencing approaches, bacteriophages were initially classified primarily based on their morphology, specifically by their head shape and tail structure as observed by transmission electron microscopy(5). The first system of classification came into being in the 1960s, and bacteriophages were grouped into families based on shared structural and biological properties. At the time, tailed phages made up the majority of isolated phages and were classified into three families based on their tail structure: *Myoviridae* (with long contractile tails), *Siphoviridae* (with long non-contractile tails), and *Podoviridae* (with short tails) within the order *Caudovirales* (6).

Recently there have been concerted efforts to provide a universal viral taxonomy across all viruses including bacteriophages and viruses of other organisms, and establish principles enabling such an approach. The first principle is that taxa should be monophyletic - that share a single common ancestor (7). As sequencing technologies have developed it has become possible to infer the evolutionary history of bacteriophages based on conserved hallmark genes such as the large terminase subunit (*terL*) or entire genomes (e.g. tBLASTx) (8). Unsurprisingly, it became apparent that the genetic diversity of phages goes far beyond the observed morphological diversity (9–11). Several studies showed that while certain morphological features might be conserved within lineages of phages, the genetic and evolutionary relationships amongst phages is significantly more complex (12). Phages with similar morphologies can have considerable genetic differences and belong to different evolutionary lineages and thus are not monophyletic (12,13), violating the first principle that taxa of viruses should represent monophyletic groups (7).

With further advances in sequencing technology and rapidly decreasing costs, there have been increasing reports that have highlighted the incongruence of morphological based taxonomy (12–14). This has stimulated a move towards genomic based classification that aims to enable a universal taxonomy for all viruses, including bacteriophages. This has resulted in the morphological classification being abolished and adoption of a binomial naming system has been adopted (15,16). The ICTV now utilises a 15-rank taxonomic framework, spanning from realm down to the basal rank of species. Each taxonomic rank, with the exception of species, has a specific suffix to allow the identification of the rank: realm (*viria*), subrealm (*vira*), kingdom (*virae*), subkingdom (*virites*), phylum (*viricota*), subphylum (*viricotina*), class (*viricetes*), subclass (*viricetidae*), order (*virales*), suborder (*virineae*), family (viridae), subfamily (*virinae*) and genus (*virus*).

As a result of the abolishment of morphology-based taxa, the iconic families *Myoviridae, Siphoviridae* and *Podoviridae* with the order *Caudovirales* have now been removed (17). While these families are no longer formal taxa, the classic morphological descriptions of podovirus, myovirus and siphovirus are still maintained, providing context to the historical literature (17) since the majority of isolated tailed bacteriophages were classified into these families *(18)*. The genera within the former class *Caudovirales* have been moved into new recently created families or remain as floating genera, within the order *Caudoviricetes*, allowing for the creation of new families and orders. The creation of new viral families can be a time consuming process that requires large scale genomic analyses to identify orthologous genes that are shared across the proposed monophyletic family (19). The creation of taxa at the level of a family and above is not easily automated and requires substantial manual curation and effort. In contrast, classification at the genus and species level is based upon average nucleotide identity (ANI) and presents the opportunity for automation to substantially speed up the process.

There are now very clear guidelines for placement of bacteriophages into genera and species (17,19). The dsDNA bacteriophages with an average nucleotide identity (ANI) ≥95% are considered the same species, and bacteriophages with an ANI ≥ 70% over 100% of the genome are considered to be within the same genus (16,19). There are a number of tools available to calculate or approximate ANI. The most simplistic is BLASTn, normalised for both the identity of the alignment and the length of the alignment to the total genome length (19). A more advanced approach and now recommended by the ICTV is to normalise for genome length and high-scoring segment pairs (HSP) from the results of BLASTn. This approach has been implemented in the Virus Intergenomic Distance Calculator (VIRIDIC) (20). VIRIDIC allows for the comparison of multiple bacteriophage genomes and produces both a graphical output and similarity matrix of intergenomic similarity. VIRIDIC is available via a web interface or a downloadable singularity distribution (20) and has become a widely-used tool in bacteriophage genome classification.

Despite the number of tools that are available to calculate the similarity between phage genomes, the process of assigning taxonomy to a newly sequenced phage genome is a non-trivial task for those not familiar with command line based tools. Furthermore, the decreasing costs of sequencing and the resurgence of bacteriophage research is resulting in the rapid expansion of the number of complete bacteriophage genomes in the INSDC that require classification (21). For classification to keep pace and to enable the Bacterial Viruses Subcommittee of the ICTV to focus on classification at higher taxonomic ranks, there is a clear need for fully automated classification of bacteriophage genomes at the levels of genus and species.

The steps required for bacteriophage genome classification are 1) identify the closest relatives of a newly sequenced bacteriophage, 2) calculation of genomic distance (ANI) compared to these relatives, 3) identify currently classified ICTV bacteriophages, and 4) determine the similarity of a newly isolated bacteriophage against ICTV classified bacteriophages. While there are tools for many of the steps, they are not integrated and data are held in multiple databases. For instance, comparison against all known bacteriophage genomes is easily done through the NCBI web blast interface (22) or INPHARED database (21). Genomic similarity can be calculated by VIRIDIC (20) via a web interface or the command line. A list of currently classified genomes is available from the ICTV website via the Virus Metadata Resource (VMR). However, without familiarity with programming, linking currently classified phages that are listed in the VMR to those available in Genbank and importing into VIRIDIC is a time consuming and laborious task that involves manipulation of data in multiple formats.

Here we sought to develop a high-throughput and easy to use system that enables the rapid classification of dsDNA bacteriophages to the genus and species level, and which scales with increasing volume of data. We present a workflow that takes a bacteriophage genome as an input and determines if the bacteriophage is a representative of any currently defined genera or species. The process removes the need to manually cross check against multiple databases, upload data to multiple websites or the ability to write scripts to automate the process. We have developed the tool taxmyPHAGE which is available as both a standalone version via conda and pip, and as a web-interface at taxmyphage.ku.dk.

## Materials and Methods

### An overview of the workflow is provided in Figure 1

We created a mash database of bacteriophage genomes that have been classified by the ICTV, sketching each genome with 5000 sketches, using a sketch size of, -s 5000 (23). The mash database can be updated with the yearly release of the ICTV Virus Metadata Resource which contains details of all classified virus genomes. The initial search against the mash database allows for rapid identification of genomes similar to the query sequence. The taxonomy of the hits identified is extracted from the ICTV VMR and then all genomes comprising those genera are extracted. The genus information is then utilised to construct a subset of genomes that the query sequence will be compared against in more detail. For instance, if the top hits from mash identified similarity to nine phages in the genera *Bristolvirus* and one phage in the genus *Bellamyvirus*, all phage genomes within these two genera are extracted and combined with the query genome for further analysis. We re-implemented in python the VIRIDIC algorithm to calculate intergenomic genomic similarities, that takes into account genome length along with query coverage to calculate average nucleotide identity (20). Using python provided considerable speed up compared to the R implementation and allowed us to scale with increasing volumes of data. Genomes are then clustered at 70% and 95% ANI to meet ICTV guidelines for the demarcation of genera and species. The query genome is then compared against these clusters to determine if 1) it is a representative of an existing species, 2) is a new species within an existing genus, 3) represents a new species within a new genus and 4) identifies if current ICTV taxonomy is incongruent with the current genomic demarcation criteria. The output provides the user with an indication of the current taxonomy. The web version is restricted to one genome at a time whereas the command line interface takes an multi-fasta input and will process each fasta entry as an individual genome.

**Figure 1.**
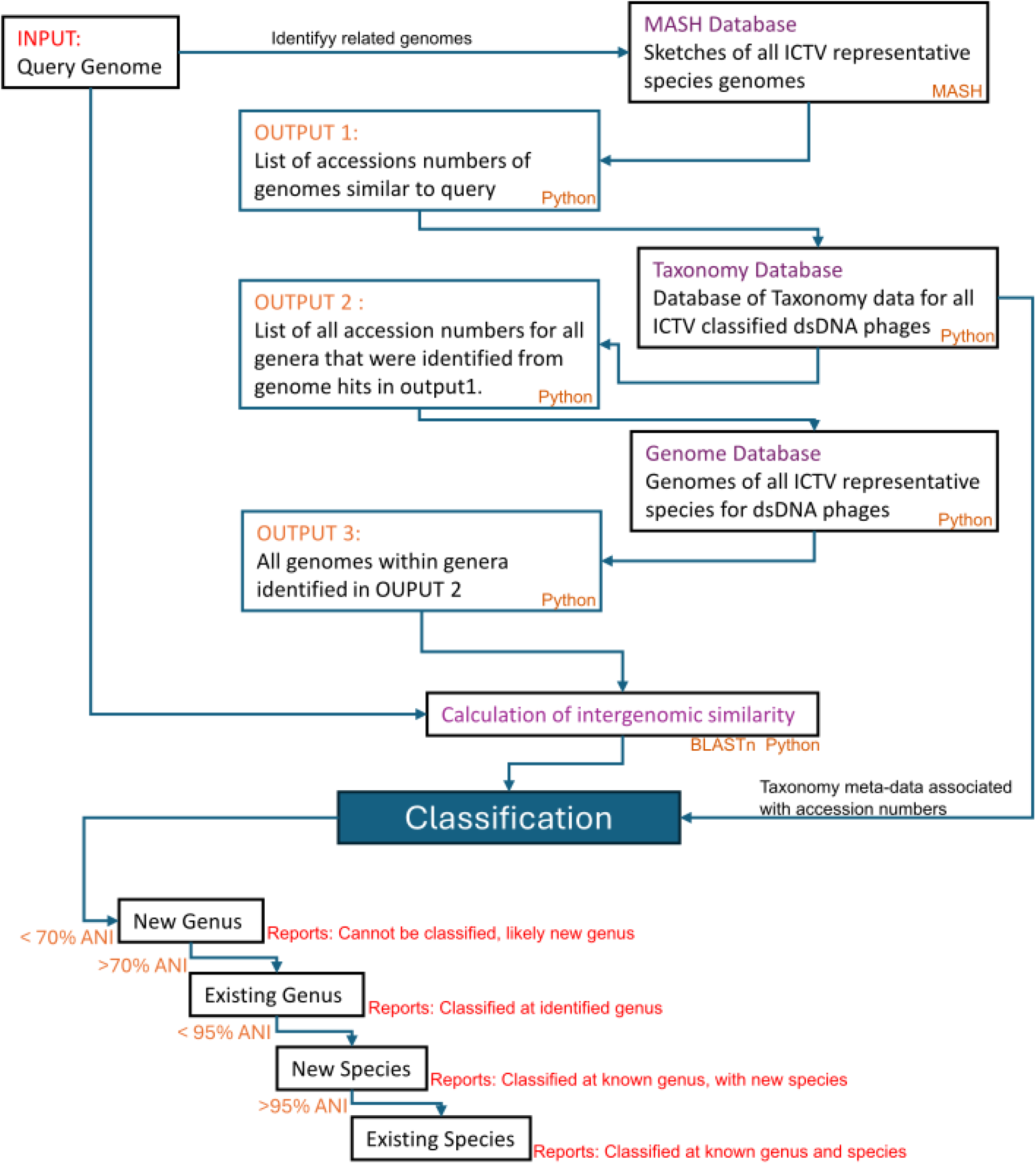
Overview of the classification process. Input is one or more query sequences in FASTA format that is compared to current ICTV classified dsDNA phage genomes, using mash. The genera of the resultant top hits are used to identify the unique genera the query is similar to. All genomes within these genera are extracted and compared to the query sequence to calculate genomic similarities. The results of genomic similarity are used to classify the query sequence into a new genus (<70% ANI), a current existing genus (≥70%), a new species (≥ 70 ANI < 95%) or current species (>95% ANI).

Benchmarking was carried out on a cloud notebook CLIMB-BIG DATA server, with Intel Xeon Processors (Skylake Model 85), with 16 threads used. To test our approach we have utilised the delay taken from when taxonomy proposals are submitted to the ICTV to the time taken for the latest virus metadata resource (VMR) to be ratified and released. We utilised the VMR_MSL38_v1.xlsx, released on 04/25/2023 to test a set of bacteriophage taxonomy proposals that were submitted to the ICTV Bacterial Viruses Subcommittee in March 2023 and later ratified by the Executive Committee in August 2023.

## Results

We developed a single workflow for the classification of dsDNA phages genomes to the genus and species level, that is available as a standalone python script available via pipy, conda, github or can be accessed via a web interface. We tested representative species from ten different genera, classification for all 10 genomes was correct. The time taken to classify a genome was dependent on the number of existing genomes within a genus and the number of closely related genera identified in initial searches of the mash database (Table 1). For instance, there are only nine species in the genus *Pseudotevenvirus*, however, the initial rapid mash searching will identify other closely related genera in the *Straboviridae* (Table 1). Genomes from all these genera are processed in the more computationally expensive blastn analysis, allowing the genus and species to be resolved for the submitted genome(s). Time for blastn analysis is dependent on both the number of genomes and the size of the genomes. Despite this, it was still possible to rapidly classify a genome sequence to the species level and provide supporting figures in less than 30 minutes for all genomes tested, a substantial time saving on all current methods that require interacting with multiple different databases or tools.

**Table 1.** Benchmarking of time taken to classify a genome. The number of genomes assigned to a genus is from the VMR v 38. The number of identified genera is from the initial MASH searching prior to the more accurate BLASTn analysis.

| Genus | Number of Genomes in assigned genus | Time (h:m:s) | Number of genera identified by MASH |
| --- | --- | --- | --- |
| *Cheoctovirus* | 96 | 00:07:44 | 1 |
| *Tequatrovirus* | 83 | 00:26:19 | 2 |
| *Peduovirus* | 27 | 00:00:23 | 1 |
| *Warwickvirus* | 18 | 00:00:18 | 2 |
| *Pseudotevenvirus* | 9 | 0:01:15 | 2 |
| *Lillamyvirus* | 6 | 0:00:12 | 3 |
| *Kablunavirus* | 3 | 0:00:27 | 3 |
| *Changmaivirus* | 2 | 0:00:17 | 1 |
| *Stompvirus* | 1 | 0:00:16 | 1 |

### Classification of new genomes

To test the accuracy of taxmyphage with new genomes, we utilised a set of 734 genomes that had been submitted for classification, but were pending approval by the ICTV executive committee and as such were not already in our mash database. The data included examples of genomes in entirely new genera, which taxmyphage will not be able to name, but can predict the genome to be representative of a new genus and species. Using this approach allowed us to test whether taxmyphage is able to assign phages to the correct genus and identify new species. As taxonomy is not static and continuously updated as genomic space is expanded, there data contains pending data of existing species/genera that are being reclassified into new taxa.

Using the command line version that allows multiple genomes to be classified from one input file, 734 genomes were classified in less than 48 hours. For 125 genomes that are pending approval into new genera and species, tamxyphage correctly identified these as representatives of new genera and species. The genus classification was congruent with the pending ICTV taxonomy for 96.7% (560/579) of the genomes tested (Supplementary table 1). Those genomes that differed were examined in more detail. Five genera account for disagreement in taxonomy, these were; *Warwickvirus* (1), *Xooduovirus* (1),*Otagovirus* (3),*Beetrevirus* (6), and *Jedunavirus* (8). For the genera *Otagovirus, Beetrevirus* and *Jedunavirus*, within the current classification system there are multiple genomes that are <70% ANI to other genomes classified in the same genus (Figure 2). Thus, taxmyphage was able to identify incongruence in the current classification system with a 70% threshold for a genus that led to misclassification of genomes (Figure 2a). For genomes in the genera *Warwickvirus* and *Xooduovirus*, the results of taxmyphage indicated they had >70% ANI to genomes in these genera, but only just at 71.3% and 70.5 %. If other tools are used to calculate ANI rather than VIRIDIC algorithm as suggested by the ICTV, then values <70% can be obtained which would result in these species incorrectly being excluded from these genera.

**Figure 2.**
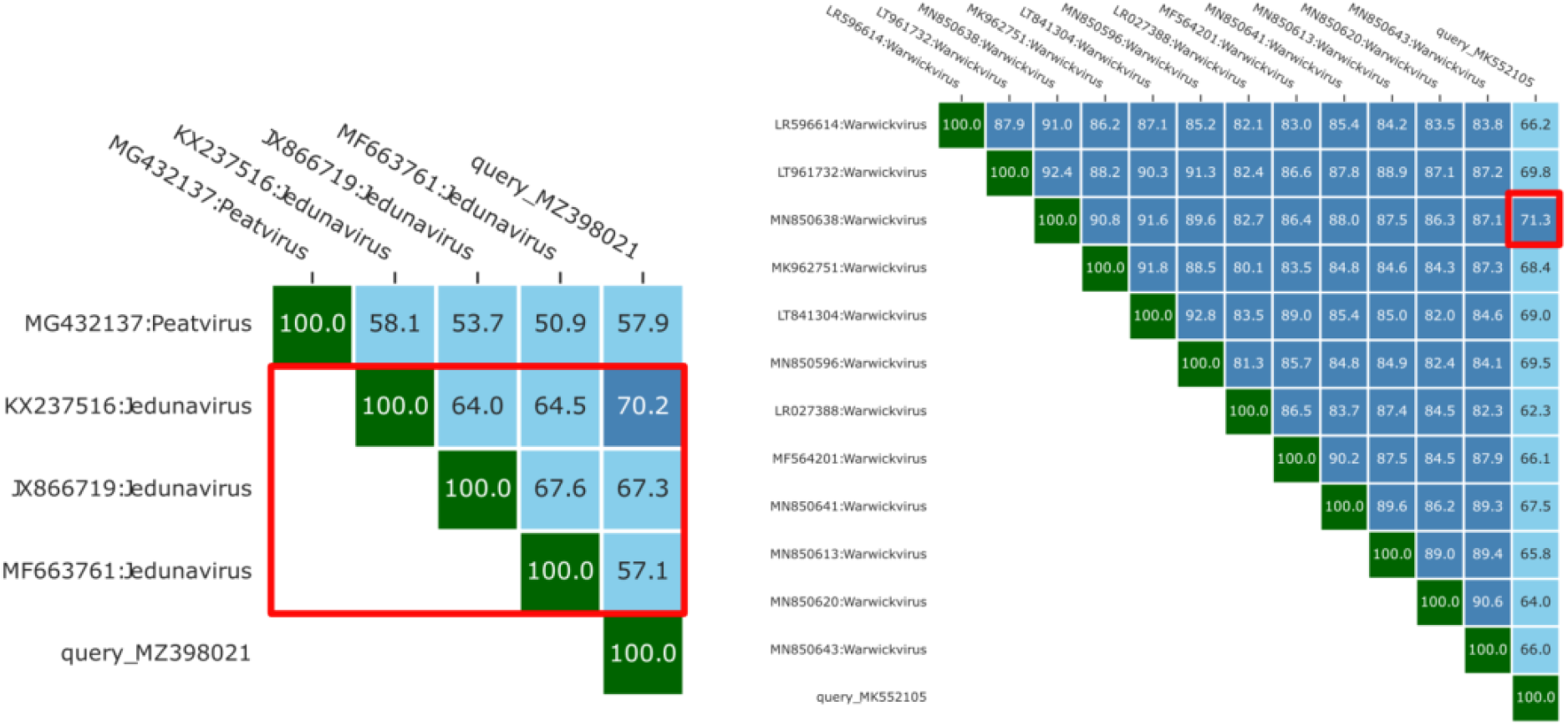
Classification of genomes MZ398021 and MK552105. a) top right matrix of genomic similarity of phage genome MZ398021 with other phages in the *Jedunavirus*. b) top right matrix of genomic similarity of phage genome MK552105 with other phages in the genus *Warwickvirus*

Within the pending ICTV classification data tested 505 new species were proposed and taxmyphage was congruent with 98.5% (492/505) of these, correctly stating the query was representative of a new species. In the other 13 cases, the phage genomes were assigned to an existing species, necessitating further detailed examination. In all 13 cases taxmyphage made an assignment to an existing species because the genome was between 95-96 % similar to an existing species. Again these differences between the pending taxonomy and results from taxmyphage may result from the multiple different methods that can be used to calculate ANI, where the difference between 94.9 and 95.1 is small but can influence taxonomy. It is noteworthy that if all these pending changes are accepted by ICTV, the data would be incorporated into the taxmyphage database and genomes would be correctly assigned to these taxa.

## Discussion

With the resurgence in bacteriophage research due to their potential as therapeutic and biocontrol agents, increasing numbers of bacteriophage genomes are being sequenced (21). In parallel, the move to a unified genome-based taxonomy requires the development of easy to use tools to enable the rapid and consistent classification of dsDNA phages. Taxmyphage now provides all generators of phage genomes the ability to classify their phages, such that phage genome sequencing and classification can be democratised and not the domain of a select few. The increase in bacteriophage genomes is exemplified by the ∼ 6500 genomes released in Genbank between March 2023 and March 2024. As of April 2023, ∼4500 bacteriophage species have been classified by the ICTV. Compared to the INPHARED database, which now contains 28,000 sequence records, there is a clear requirement for the development of rapid, easy to use tools capable of scaling with increasing amounts of data for the classification of bacteriophages to address the large backlog of bacteriophage genomes that remain without taxonomy.

Taxmyphage builds on the algorithm developed in VIRIDIC (20), resulting in a substantial increase in speed when implementing the algorithm in python that allows for larger datasets to be analysed, overcoming the bottleneck associated with VIRIDIC. Furthermore, unlike other tools such as VIRIDIC (20) and VICTOR (24), it does not require any *a priori* knowledge of the closest relatives to correctly identify the taxonomy of a query sequence. In summary, taxmyphage provides a one stop solution for the classification of bacteriophages at the lower taxonomic ranks of genus and species. The web interface allows users with no experience of bioinformatics to rapidly and accurately classify their phage genomes. The command line version allows more advanced users to incorporate the process into existing workflows. As such taxmyphage has the potential to substantially increase the rate and number of bacteriophage genomes that are classified at the levels of genus and species.

## Acknowledgements

For the purpose of open access, the author has applied a Creative Commons Attribution license (CC BY) to any Author Accepted Manuscript version arising from this submission. A.M is funded by MRC (MR/L015080/1 and MR/T030062/1). Bioinformatics analysis was carried out on infrastructure provided by MRC-CLIMB (MR/L015080/1).

